# Identifying Clusters in Graph Representations of Genomes

**DOI:** 10.1101/2023.07.20.549917

**Authors:** Eva Herencsárová, Broňa Brejová

## Abstract

In many bioinformatics applications the task is to identify biologically significant locations in an individual genome. In our work, we are interested in finding high-density clusters of such biologically meaningful locations in a graph representation of a pangenome, which is a collection of related genomes. Different formulations of finding such clusters were previously studied for sequences. In this work, we study an extension of this problem for graphs, which we formalize as finding a set of vertex-disjoint paths with a maximum score in a weighted directed graph. We provide a linear-time algorithm for a special class of graphs corresponding to elastic-degenerate strings, one of pangenome representations. We also provide a fixed-parameter tractable algorithm for directed acyclic graphs with a special path decomposition of a limited width.

## 1. Introduction

The rapid decreases in the cost of genome sequences led to a shift in genomics and bioinformatics from analyzing a single representative genome per species to analyzing genomes of many individuals. A collection of related genomes analyzed jointly is called a *pangenome* [1]. Pangenomes are often represented as graphs, in which nodes correspond to parts of the sequences and edges to adjacencies between these sequences observed in at least one of the studied genomes [2, 3].

Introduction of pangenome graphs gave rise to a need to extend many bioinformatics algorithms from working with single sequences (strings) to graphs representing a family of related sequences. In this work, we introduce algorithms that identify clusters of biologically meaningful positions in such pangenome graphs. In many areas of bioinformatics, one can identify genome positions having some biological function or property and then search for dense clusters of such positions. The simplest examples are based on sequence content, such as looking for GC-rich regions (regions with high density of bases C and G) [4] or CpG islands (regions with high density of C followed by G) [5]. Such areas are often associated with functional elements such as genes or regulatory regions [6, 7]. A more complex example is looking for clusters of motifs representing transcription factor binding sites [8]. We can also identify positions of mutations within or between species and look for conserved regions lacking such mutations [9] or regions with a high density of mutations arising for example from horizontal sequence transfer [10]. All of these examples involve identifying individual bases with some biological property and then looking for groups of such bases located close together.

One possible formalization of locating such clusters in a single DNA sequence is to assign a score to each base which is positive for bases with the property of interest and negative for other bases, and then look of high-scoring intervals in the resulting sequence of scores. Miklós Csűrös [4] formulated this approach as looking for a set of disjoint intervals with maximum sum of scores, where the user either restricts the number of intervals to *k* or assigns a negative penalty to each interval in the output set. The latter problem can be solved in linear time by a simple dynamic programming algorithm, and will form the basis of the approach outlined in this article.

Namely, we generalize the maximum-scoring segment set problem [4] from sequences of scores to weighted directed graphs representing pangenomes. The weights of individual edges represent scores of bases in a pangenome. In a sequence, a cluster is typically defined as a contiguous segment (interval). In the graph extension, one can consider various definitions of the concept of a segment, such as a connected induced subgraph. However, we have decided to look for clusters defined as paths in the graph, as each path corresponds to a single sequence (either one of the constituent genomes of the pangenome or a combination of multiple such genomes). This gives rise to the maximum-score disjoint paths problem defined in the next section. In section 3 we provide a linear-time algorithm for a special class of graphs corresponding to elastic-degenerate strings [11]. In section 4 we give an algorithm for general directed acyclic graphs. The complexity of this algorithm is exponential in a parameter of a special path decomposition of the graph, but linear in the overall size of the graph.

### 2. Notation and problem definition

In this work, we will consider a weighted directed graph *G* with vertex set *V*, edge set *E* ⊆*V* ^2^ and weight function *w* : *V* →ℝ. We will first introduce graph terminology and notation used in this work. For each edge (*u, v*) ∈ *E* we call *u* a predecessor of *v* and *v* a successor of *u*. The set of all predecessors of *v* is denoted *N* ^−^(*v*). The subgraph of *G* induced by set *X* ⊆ *V* is the graph *G*^*′*^ = (*X, E* ∪ *X*^2^). A *path* is a sequence of distinct vertices (*v*_1_, *v*_2_, …, *v*_*n*_) such that (*v*_*i*_, *v*_*i*+1_) ∈ *E* for *i* = 1, 2, …, *n* − 1. A cycle is a path such that (*v*_*n*_, *v*_1_) ∈ *E*. If *G* does not contain a cycle, we call it a directed acyclic graph (DAG). Vertices of each DAG can be ordered topologically as *v*_1_, … *v*_*n*_ so that for each edge (*v*_*i*_, *v*_*j*_) ∈ *E* we have *i* < *j*.

We are now ready to state our problem. The goal of the *maximum-score disjoint paths* problem is for a given graph *G* and penalty *x* ∈ ℝ^+^ to find a set of vertex disjoint paths with the maximum sum of scores. The score of a single path *P* = (*v*_1_, *v*_2_, …, *v*_*n*_) is defined as 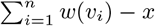. Figure 1 shows an example of the input and output for this problem.

**Figure 1:**
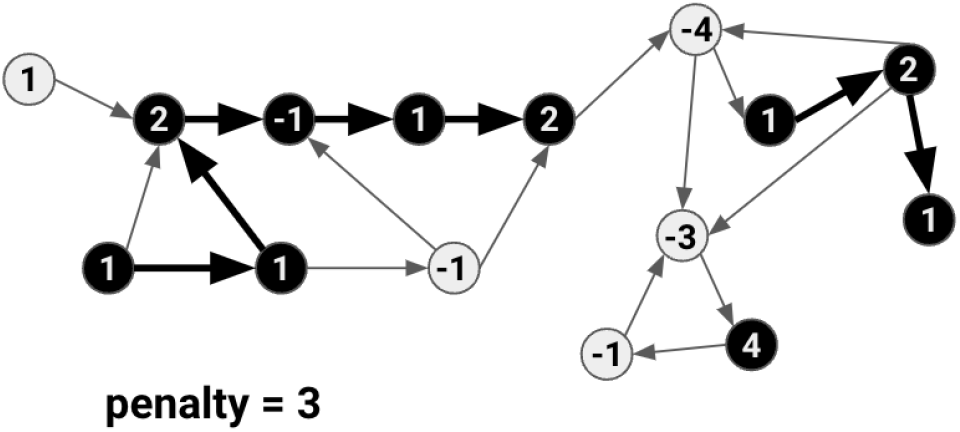
An example of a weighted directed graph and the set of paths forming the solution to the maximum-score disjoint paths problem for penalty *x* = 3. The score of this solution is (1 + 1 + 2 − 1 + 1 + 2 − *x*) + (1 + 2 + 1 − *x*) + (4 − *x*) = 5.

Note that the maximum-score disjoint paths problem is NP-hard for arbitrary weighted directed graphs. The NP-hardness can be easily proved by a reduction from the Hamiltonian path problem. If we set the weight of each vertex to 1 and penalty also to 1, the graph has a Hamiltonian path if and only if the maximum-score disjoint paths problem has a solution with score |*V*| −1. We will concentrate on DAGs. Our algorithms are an extension of the dynamic programming algorithm by Csűrös [4] for sequences of scores. The related problems of finding a single segment with maximum score or *k* segments in a sequence was studied by multiple authors [4, 12, 13]. A single path can also be found on a weighted tree [14, 15]. There are also algorithms for the related maximum density segment problem [16].

### 3. An algorithm for *n*-layered bubble graphs

In this section we present a linear-time algorithm based on dynamic programming for a special class of directed acyclic graphs, which we call *n*-layered bubble graphs.

#### Definition 3.1

(*b-layered bubble*). *A b*-layered bubble *is a directed acyclic graph with a start vertex s, an end vertex t and b non-empty vertex-disjoint directed paths, referred to as layers, connecting s and t*.

#### Definition 3.2

(*n-layered bubble graph*). *An n*-layered bubble graph *can be constructed by taking a sequence of vertices u*_1_, …, *u*_*k*_ *and connecting each pair of u*_*i*_ *and u*_*i*+1_ *by an edge or by a b-layered bubble with the start vertex u*_*i*_ *and the end vertex u*_*i*+1_ *and with* 2 *≤ b ≤ n*.

An example of a 3-layered bubble graph can be seen in the bottom part of Figure 2.

**Figure 2:**
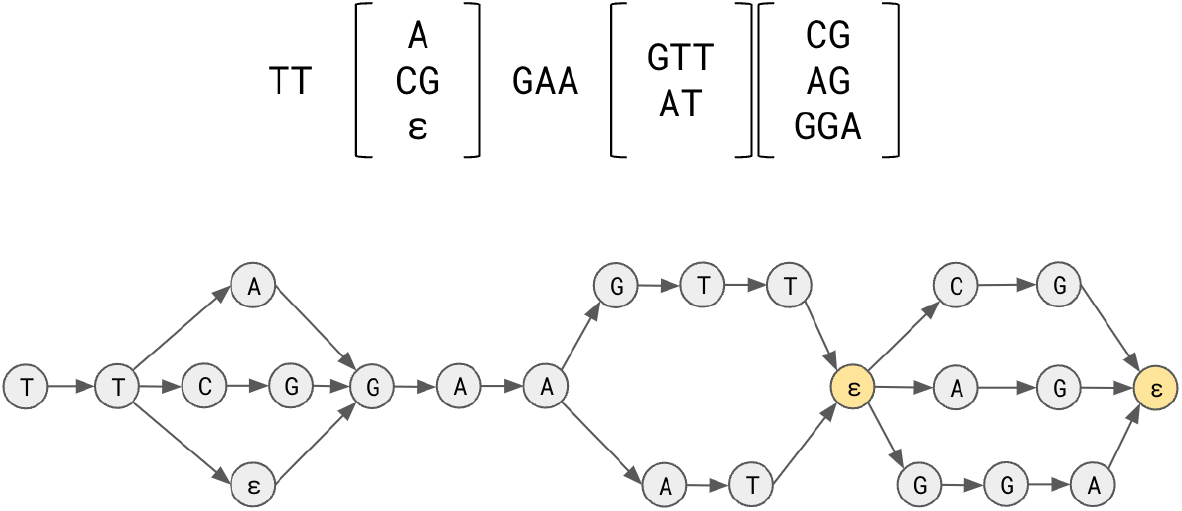
An example of an elastic-degenerate string and the corresponding 3-layered bubble graph. The bases are represented as the vertices, their adjacencies as edges. Some vertices contain an empty string denoted as *ε*.

#### Connection to elastic degenerate strings

Although the structure of the *n*-layered bubble graphs is very simple, they correspond to a well-studied representation of pangenomes called elastic-degenerate strings (EDSs) [11]. An EDS is a string containing *elastic-degenerate symbols*. An elastic degenerate symbol is defined as a set of strings, potentially of different lengths. Thus the EDS represents a set of strings, each obtained by choosing one of the strings from each elastic degenerate symbol and concatenating them.

An EDS with each set containing at most *n* strings can be easily converted to an *n*-layered bubble graph by replacing each elastic-degenerate symbol with a bubble, each path spelling the string one character per node. We add start and end vertices with zero weight for each bubble. Due to zero weight, they do not influence the score of the solution. If an elastic-degenerate symbol contains an empty string in its set, the path for this string will also contain an auxiliary node with zero weight. An example of a conversion of an EDS to a graph is shown in Figure 2.

#### An algorithm for a simple path

Before giving the full algorithm for *n*-layered bubble graphs, we will consider the algorithm for a simple path (*v*_1_, …, *v*_*n*_). This algorithm is very similar to the dynamic programming algorithm given by Csűrös [4] except for a slightly different meaning of the selection value *s* defined below. The algorithm fills a two-dimensional matrix *W*. For ≤ 1 ≤ *i n* and *s* ∈ {0, 1}, value *W* (*i, s*) is the score of the optimal solution using only vertices *v*_1_, …, *v*_*i*_. If *s* = 1, we further constrain the solution to include the last vertex *v*_*i*_ in one of the selected paths. If *s* = 0, we place no further constraints on the solution. Values *W* (*v*_*i*_, *s*) are computed for increasing values of *i* using the following equations:

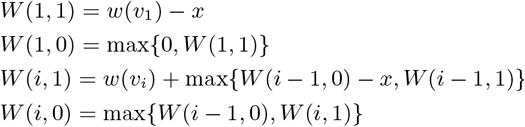

For *s* = 1, we always use vertex *v*_*i*_ with score *w*(*v*_*i*_). One option is that it starts a new path, incurring penalty of *x*. The rest of the solution will use only nodes *v*_1_, …, *v*_*i*−1_, thus having score *W* (*i* −1, 0). If vertex *v*_*i*_ continues an existing path, we instead use sub-problem *W* (*i* −1, 1) ensuring that such a path exists. For *s* = 0, we consider the case when *v*_*i*_ was used in the path, which has score *W* (*i*, 1) and the case when it was not used, which has score *W* (*i* − 1, 0).

#### An algorithm for bubble graphs

Let *G* = (*V, E*) be an *n*-layered bubble graph. We will partition its vertices into sets *N, J, L*_1_, …, *L*_*n*_ as follows. Each *b*-layered bubble in the graph consists of a start vertex, an end vertex and *b* disjoint paths *q*_1_, …, *q*_*b*_ for 2 ≤*b* ≤*n*. We will place internal vertices of each path *q*_*i*_ to set *L*_*i*_ (the ordering of the paths within the bubble is arbitrary but fixed). The end vertex of the bubble will be placed to set *J*. All remaining vertices will be placed to set *N*. Using the notation from Definition 3.2, set *N* includes vertex *u*_1_ and any vertex *u*_*i*_ which has a single predecessor. We further split each *L*_*i*_ into sets *L*_*i,first*_ and *L*_*i,later*_, where *L*_*i,first*_ contains vertices from *L*_*i*_ that do not have a predecessor in *L*_*i*_, and *L*_*i,later*_ = *L*_*i*_ \*L*_*i,first*_.

In our algorithm we will process the vertices in order *O* = *v*_1_, …, *v*_|*V* |_, which is a topological order of the graph, and in which for each bubble we first list its vertices from *L*_1_, then from *L*_2_ and so on.

Our dynamic programming algorithm fills a three-dimensional matrix of scores *W* (*i, s, l*), where *v*_*i*_ ∈ *V, s* ∈ {0, 1} is a selection value, and *ℓ* ∈ {*I, E*} is a path continuation value. Value *W* (*i, s, ℓ*) is the best score among all sets of disjoint paths within some induced sub-graph of *G* satisfying some additional properties specified below.

The induced subgraph considered in *W* (*i, s, ℓ*) is defined as follows:

- for *v*_*i*_ ∈ *N* ∪ *J* ∪ *L*_1_: the subgraph induced by *{v*_1_, …, *v*_*i*_*}*,
- for *v*_*i*_ ∈ *L*_*i*_, 2 *≤ i ≤ n* in a bubble *B*: the subgraph induced by *{v*_1_, …, *v*_*i*_*}* ∩ *ℓ*_*i*_ ∩ *B*.

Thus only the first layer of the bubble contains information about all previous vertices of the graph, whereas in the remaining layers we compute only a local score along one path of the bubble.

The selection value *s* constrains the set of paths in the same way as in the simpler algorithm above:

- *s* = 1 means *v*_*i*_ is selected in a path,
- *s* = 0 means *v*_*i*_ may or may not be selected in a path (no constraint),

The constraint imposed by the path continuation value *ℓ* depends on the type of the vertex and allows us to ensure that a path entering a bubble from its start vertex will continue in at most one layer of the bubble. For *v*_*i*_ ∈ *L*_1_:

- *ℓ* = *I*: there is no selected path which contains both the bubble’s start vertex and the subsequent vertex from *L*_*k,first*_ where *k* > 1 (no constraint on *L*_1,*first*_).
- *ℓ* = *E*: the bubble’s start vertex is selected on a path which continues with a vertex from *L*_*k,first*_ where *k* > 1.

For *v*_*i*_ ∈ *L*_*k*_, *k* > 1:

- *ℓ* = *I*: the path from the bubble’s start vertex continues on layer *L*_*k*_, i.e. both the bubble’s start vertex and the *L*_*k,first*_ vertex of the current bubble are selected;
- *ℓ* = *E*: there is no path containing both the current bubble’s start vertex and the *L*_*k,first*_ vertex of the current bubble.

For *v*_*i*_ ∈ *N* ∪ *J*, we will use only *ℓ* = *I* and do not impose any additional constraint. Value *W* (*v*_*i*_, *s, E*) is not defined and can be considered as being. −∞

To initialize the algorithm for the first node *v*_1_ in ordering *O*, we use similar formulas, as for the simpler case of a single path:

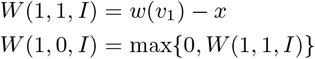

Since *v*_*i*_ ∈ *N*, we have *W* (1, 0, *E*) = *W* (1, 1, *E*) =−∞

Let us now consider some vertex *v*_*a*_ for *a* ≥ 2. We will consider several cases. If *v*_*a*_ ∉ *J*, it has a single predecessor, which we denote *v*_*p*_. The simplest case is analogous to the algorithm operating on a single path, and applies to three cases: (1) *v*_*a*_ ∈ *N* and *ℓ* = *E*, (2) *v*_*a*_ ∈ *L*_*k,later*_ for any *k* and any *ℓ* ∈ *{I, E}*, and (3) *v*_*a*_ ∈ *ℓ*_1,*first*_, *ℓ* = *I*.

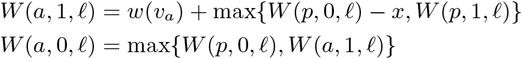

Note that the value of *ℓ* is propagated along the layers in the bubble.

The next case is *v*_*a*_ ∈ *ℓ*_1,*first*_ and *ℓ* = *E*. Value *ℓ* = *E* means that the path from the predecessor (start of the bubble) continues to some other layer of the bubble, and thus we always apply the penalty if *v*_*a*_ is included in a path. Also, the predecessor *p* is constrained to be on a path, and thus we use *W* (*p*, 1, *I*) instead of *W* (*p*, 0, *I*).

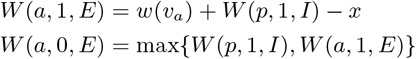

If *v*_*a*_ ∈ *L*_*k,first*_ where *k* > 1, we will not use scores computed for the predecessor, because those are propagated along the first layer. For *ℓ* = *E* we use similar formulas as for *v*_1_. For *ℓ* = *I* the path has to continue from *v*_*p*_ to *v*_*a*_, leading to more constrained formulas.

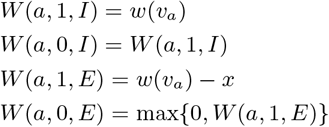

Finally, we will consider the most complex case *v*_*a*_ ∈ *J*, that is, the end vertex of a processed bubble with *b* layers. Vertex *v*_*a*_ has in this case *b* predecessors denoted here as 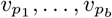, where 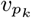 is in layer *L*_*k*_. The values stored in score matrix *W* for *p*_1_, …, *p*_*b*_ were calculated for *b* disjoint subgraphs, and thus to get the score for *a*, the algorithm has to sum up the scores for *p*_1_, …, *p*_*b*_, while ensuring that both at the start and end of the bubble the penalties for new paths are applied properly.

To ensure that the selected path from the bubble’s start vertex continues with at most one vertex *v* ∈ *L*_*k,first*_, 1 ≤*k* ≤*b*, we have to use *ℓ* = *I* for exactly one predecessor and *ℓ* = *E* for all the others. Recall that the score for *p*_1_ when *ℓ* = *I* also includes the possibility that the selected path does not continue to any of the layers from the bubble’s start vertex.

To calculate the score for *W* (*a*, 1, *I*) efficiently, three groups of sums are created and their maximum is used as *W* (*a*, 1, *I*).

The first group corresponds to the situation where we do not ensure the selection of any of the predecessors, and therefore, *v*_*a*_ starts a new path which incurs a penalty.

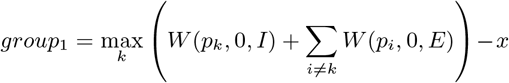

The maximum in *group*_1_ goes over *b* sums, each having path continuation value *I* for a different predecessor. The value *group*_1_ can be calculated in *O*(*b*) time. First, the sum *W* (*p*_1_, 0, *E*) + *W* (*p*_2_, 0, *E*) + + *W* (*p*_*b*_, 0, *E*) − *x* is calculated. Then the algorithm changes exactly one addend at a time from *W* (*p*_*k*_, 0, *E*) to *W* (*p*_*k*_, 0, *I*) and chooses the maximum sum.

The second group corresponds to the situation when some predecessor *v*_*pk*_ is selected (i.e. selecting *v*_*a*_ does not incur a penalty), and *ℓ* = *I* for the same predecessor *v*_*pk*_ :

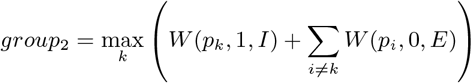

Similarly as for *group*_1_, value *group*_2_ can be calculated in *O*(*b*) time by first calculating the sum *W* (*p*_1_, 0, *E*) + *W* (*p*_2_, 0, *E*)+ +*W* (*p*_*b*_, 0, *E*), and then always changing exactly one addend at a time from *W* (*p*_*k*_, 0, *E*) to *W* (*p*_*k*_, 1, *I*) and choosing the maximum sum at the end.

The third group corresponds to the situation when some predecessor *v*_*pk*_ is selected (i.e. selecting *v*_*a*_ does not incur a penalty), and *ℓ* = *I* for another predecessor *v*_*pj*_ where *k* ≠ *j*. In this case Θ(*b*^2^) sums of length *b* need to be calculated and compared. This could be done in *O*(*b*^2^) time similarly as above. However, with some care, the maximum sum can be calculated in *O*(*b*) time as follows. The algorithm first calculates the sum *W* (*p*_1_, 0, *E*)+*W* (*p*_2_, 0, *E*)+ +*W* (*p*_*b*_, 0, *E*) as in group 2. Then it finds addends *W* (*p*_*y*_, 0, *E*) and *W* (*p*_*z*_, 0, *E*) which when replaced with *W* (*p*_*y*_, 0, *I*) and *W* (*p*_*z*_, 1, *E*) maximize the sum.

To do this, the algorithm first finds *p*_*c*_ and *p*_*d*_ (*c* ≠ *d*) for which the difference *W* (*p*_*y*_, 0, *I*) − *W* (*p*_*y*_, 0, *E*) is the largest and second largest, respectively. Next, it finds *p*_*e*_ and *p*_*f*_ (*e* ≠ *f*) for which the difference *W* (*p*_*z*_, 1, *E*) − *W* (*p*_*z*_, 0, *E*) is the largest and second largest, respectively. Both these computations can be done in *O*(*b*) time. Finally we use these values to assemble the final value for group 3. If *pc* ≠ *p*_*e*_, then the addend *W* (*p*_*c*_, 0, *E*) is replaced with *W* (*p*_*c*_, 0, *I*), and *W* (*p*_*e*_, 0, *E*) with *W* (*p*_*e*_, 1, *E*). If *p*_*c*_ = *p*_*e*_, then one of the addends is replaced by *W* (*p*_*d*_, 0, *I*) or *W* (*p*_*f*_, 1, *E*) instead, whichever results in a larger sum.

Finally, the value of *W* (*a*, 1, *I*) is derived in the following way:

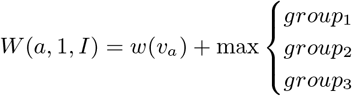

To compute *W* (*a*, 0, *I*), we take the maximum of *W* (*a*, 1, *I*) representing the case that *v*_*a*_ is selected and the value *group*_4_ representing the case that *v*_*a*_ is not selected. The value of group 4 is computed similarly as group 1, except that the penalty term − *x* is not applied. For *v*_*a*_ ∈ *J*, value *W* (*a*, 0, *E*) and *W* (*a*, 1, *E*) are not defined and can be considered as being. −∞

Once the algorithm fills in the entire matrix *W*, the overall score can be found in *W* (|*V* |, 0, *I*). Note the the last vertex *v*_|*V* |_ belongs to *N* ∪ *J*, and thus value for *ℓ* = *E* is undefined. To reconstruct the set of paths leading to the optimal score, we can store for each value of matrix *W* which case was used to obtain it and the follow these values from *W* (|*V*|, 0, *I*) all the way to *W* (1, ?, *I*).

Regarding the time complexity of the algorithm, calculating the scores for each vertex outside of *J* is done in *O*(1) time. Calculating the scores for a vertex *v*_*a*_ ∈ *J* with indegree *b* takes *O*(*b*) time, but this can be amortized among the *b* predecessors of *v*_*a*_, each of which has indegree 1. Therefore both the time and space complexity of the algorithm is *O*(|*V* |).

### 4. An algorithm for general DAGs

In the previous section, we described an algorithm for the maximum-score disjoint paths problem on *n*-layered bubble graphs. Although such graphs can provide a representation of a pangenome, their power is limited. In this section we provide a fixed-parameter tractable algorithm for a general DAG, which can solve the problem in time (2^*w*^ ·*w* ·|*V*|) if it is provided with a special directed path decomposition with the width bounded by parameter *w*. In this section we first define this decomposition and then describe the algorithm.

**Definitions**. We define the decomposition and its width in the next definition, see also example in Figure 3.

**Figure 3:**
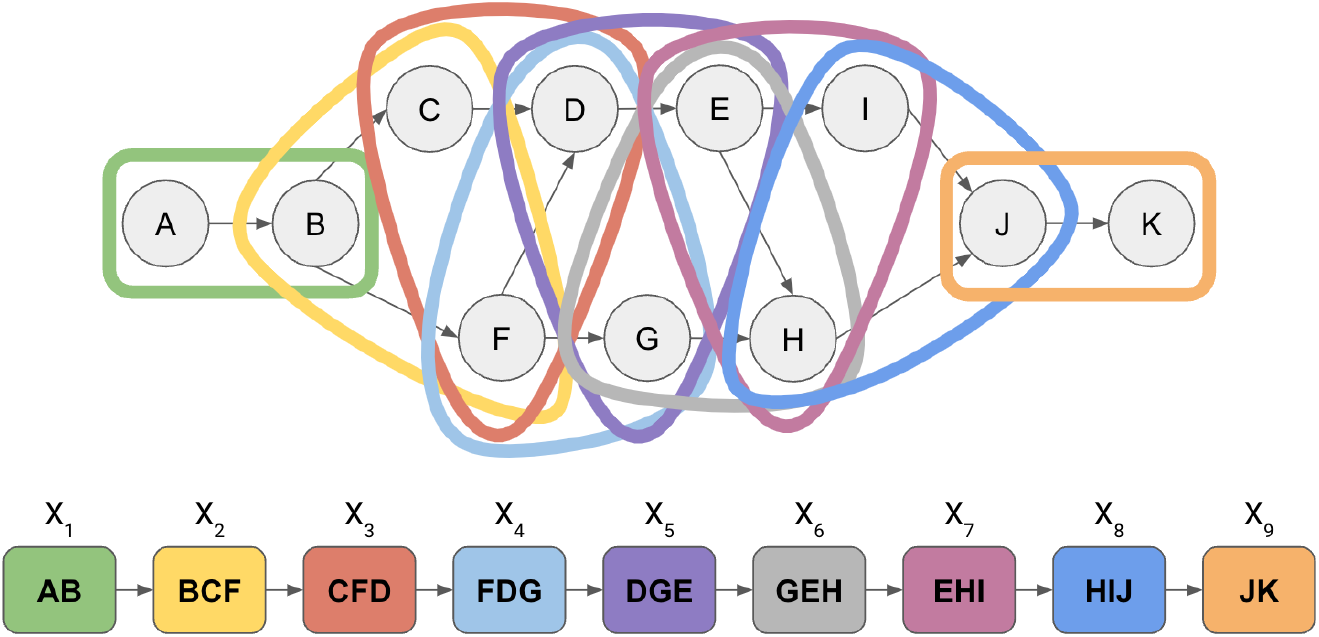
An example of a directed path decomposition with width 2 according to Definition 4.1.

#### Definition 4.1

(*Directed path decomposition*). *Let G* = (*V, E*) *be a directed graph. The* directed path decomposition *of G is a sequence of subsets* (*X*_1_, …, *X*_*n*_) *of V (we refer to them as bags of vertices), with three properties:*

i. *For each edge* (*u, v*) ∈ *E, there exists an i ∈{*1, …, *n} such that both u and v belong to bag X*_*i*_.
ii. *For every three indices* 1 *≤ i ≤ j ≤ k ≤ n, X*_*i*_ ∩ *X*_*k*_ *⊆ X*_*j*_.
iii. *For each edge* (*u, v*) ∈ *E if v* ∈ *X*_*j*_ *then there exists a bag X*_*i*_ *containing u where i ≤ j*.

*The* width *of the path decomposition is k* = max_*i*∈ *{*1,…,*n}*_ |*X*_*i*_| − 1.

One can also define the directed pathwidth of graph *G* as the minimum value *w* such that *G* has a path decomposition with width *w*.

Our definitions of a directed path decomposition and a directed pathwidth are extensions of the well-studied path decomposition for undirected graphs [17]. The path decomposition of graph *G* can be interpreted as a *thickened* path graph. The path width is a value describing that how much this path is thickened to get *G*. To adapt the undirected path decomposition for our purposes, we added the third condition. It allows the algorithm to process bags in order and ensure that predecessors of each node are already processed when we process the first bag with this node.

A different path decomposition was previously studied [18, 19], which omits the first condition and uses a less strict version of the third condition as follows: “For each edge (*u, v*) ∈ *E* there exists *i* ≤ *j* such *u* ∈ *V*_*i*_ and *v* ∈ *V*_*j*_”. However, such a relaxed definition does not seem to lead to an efficient algorithm for our problem.

The following lemma shows a useful property of a directed path decomposition.

#### Lemma 4.1.

*Let G* = (*V, E*) *be a directed graph and P* = (*X*_1_, …, *X*_*n*_) *its directed path decomposition. Assume X*_*i*_ *is the bag where vertex v appears for the first time in P, i*.*e. v* ∈ *X*_*i*_ *and v* ∉ *X*_*j*_ *where j* < *i. Then bag X*_*i*_ *contains all predecessors of v*.

*Proof*. From (*iii*) in Definition 4.1 we know that all predecessors of *v* have to be in bag *X*_*i*_ or bag *X*_*h*_ where *h* < *i*:

- If the predecessor vertex *p* ∈ *X*_*i*_, then the statement holds for *p*.
- If *p* ∈ *X*_*h*_, where *h* < *i*: Since *X*_*i*_ is the bag where *v* appears for the first time in *P*, based on (*i*) in Definition 4.1 there exists a bag *X*_*k*_, where *k ≥ i*, containing vertices *p* and *v*. Based on (*ii*) in Definition 4.1 *X*_*i*_ contains *p* ∈ *X*_*h*_ ∩ *X*_*k*_.

#### Corollary 4.1.

*The pathwidth of a directed graph G is at least the maximum indegree of G, where the indegree of a vertex v is the number of v’s predecessors*.

In our algorithm, we will use a special form of the directed path decomposition, in which a single new node is added in each bag. Below we define it formally and show that any directed path decomposition can be efficiently converted into this form without increasing the width.

#### Definition 4.2

(*Incremental path decomposition*). *Let G* = (*V, E*) *be a DAG and P* = (*X*_1_, …, *X*_*n*_) *its directed path decomposition. We consider X*_0_ = *∅. We call P an* incremental path decomposition *if* |*X*_*i*_ *∖ X*_*i*−1_| = 1 *for* 1 ≤ *i* ≤ *n. The vertex in X*_*i*_ *∖X*_*i*−1_ *is called the* incremental vertex.

Note that an incremental path decomposition of a DAG *G* = (*V, E*) consists of exactly |*V*| bags, as exactly one vertex is added in each bag and each vertex needs to be added exactly once.

#### Lemma 4.2.

*Let G* = (*V, E*) *be a DAG and P* = (*X*_1_, …, *X*_*n*_) *its directed path decomposition of width w. It can be converted to an incremental path decomposition for G with a width at most w in O*(*w ·* |*V* |) *time*.

*Proof*. Let us assume that |*X*_*i*_ *∖ X*_*i*−1_| = *k*. If *k* = 0 then *X*_*i*_ ⊆ *X*_*i*−1_ and therefore *X*_*i*_ can be left out of the path decomposition without breaking properties (*i*), (*ii*) and (*iii*) from Definition 4.1. If *k* > 1, we create a path decomposition *P* ^*′*^ = *X*_1_, … *X*_*i*−1_, *Y, X*_*i*_, … *X*_*n*_ where |*Y∖ X*_*i*−1_ |= 1 and |*X*_*i*_ *∖Y*| = *k* − 1. By repeating these steps, we get an incremental path decomposition.

To construct *Y*, we consider a topological order of vertices in *G* and select the vertex *v* which is the first in this topological order among vertices in *X*_*i*_ *∖ X*_*i*−1_. This means that *v* has no predecessor in *X*_*i*_ *∖ X*_*i*−1_. Bag *Y* is constructed as *Y* = (*X*_*i*−1_ ∩ *X*_*i*_) ∪ {*v*}. Clearly, decomposition *P* ^*′*^ satisfies all properties from Definition 4.1. Also notice that |*Y*| ≤ |*X*_*i*_|, i.e. the width of the path decomposition was not increased.

Finally, set *Y* can be constructed in *O*(*w*) time, and as repeat this process at most |*V*| times, the total running time if *O*(*w* · |*V*|). The topological order can be computed in *O*(|*V*| + |*E*|) time. Note that |*E*| ≤ *w* · |*V*| as each vertex has at most *w* incoming edges.

#### An algorithm that uses an incremental decomposition

We now describe an algorithm for solving the maximum-score disjoint paths problem for a DAG *G* = (*V, E*) and penalty *x*. The input to the algorithm is an incremental path decomposition *P* = (*X*_1_, …, *X*_*n*_) of *G*. The algorithm runs in *O*(2^*w*^ ·*w*· |*V*|) time where *w* is the width of *P*. Let *G*_*i*_ be the subgraph of *G* induced by vertices in *X*_1_ ∪ *·* ∪ *X*_*i*_.

The algorithm processes individual bags in the decomposition one at a time. When processing bag *X*_*i*_ the algorithm computes maximum scores of solutions in the subgraph *G*_*i*_. In the first algorithm, we have considered for each ending vertex *v*_*i*_ solutions for different settings of binary variables *s* and *ℓ*. Here we will consider 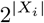 different solutions, each corresponding to a different subset *A* ⊆ *X*_*i*_. We will these subsets *configurations*. Configuration *A* determines which vertices from *X*_*i*_ are the last vertices in individual paths contained in a solution of the problem. The algorithm thus computes a score matrix *W* (*i, A*) which contains the score of the best set of paths within *G*_*i*_ such that if *B* is the set of last vertices of these paths, then *A* = *B* ∩ *X*_*i*_.

Let us assume now that the scores for *X*_*i*−1_ are already known, and we want to calculate scores for *X*_*i*_. Let *v*_*i*_ be the incremental vertex of *X*_*i*_. Consider a configuration *A* ⊆ *X*_*i*_. We will consider two cases.

First, if *v*_*i*_ /∈ *A*, then the incremental vertex is not used in any path because it is not the last vertex of any path, and it cannot be in the middle of a path, as it does not have any successors in *G*_*i*_. Therefore we copy some score computed for *X*_*i*−1_ to *W* (*i, A*). However, we need to consider multiple configurations for *X*_*i*−1_ as there can be multiple vertices in *X*_*i*−1_ which are not part of *X*_*i*_. These vertices can be part of a configuration for *X*_*i*−1_ but are no longer relevant for *X*_*i*_. To this end, for each configuration *A* of *X*_*i*_ we will define set *p*(*i, A*) of configurations of *X*_*i*−1_ that agree with *A* on the vertices shared between *X*_*i*−1_ a *X*_*i*_. Formally,

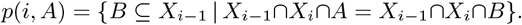

Score *W* (*i, A*) can then computed as follows:

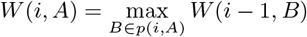

In the second case, *v*_*i*_ ∈ *A*. The path containing *v*_*i*_ can be either a single vertex, in which case apply penalty *x*, or *v*_*i*_ can follow some vertex *u* ∈ *X*_*i*−1_. In that case *u* must be in the configuration for *X*_*i*−1_, because it was the last vertex before addition of *v*_*i*_. But it is not in the configuration for *X*_*i*_, because it is now followed by *v*_*i*_. We consider all possibilities for predecessor *u* of *v*_*i*_ which is not in *A* and for configuration *B* for *X*_*i*−1_ which contains *u* but otherwise agrees with *A* on the vertices shared between *X*_*i*−1_ a *X*_*i*_. Note that all predecessors of *v*_*i*_ are in both *X*_*i*_ (according to the definition of a directed path decomposition) and *X*_*i*−1_ (because only a single vertex is added to *X*_*i*_).

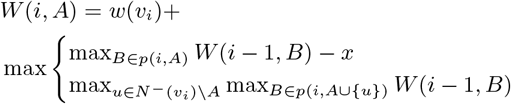

To initialize the algorithm, we set *W* (0, *∅*) = 0. The final score is the maximum of *W* (|*V* |, *A*) among all configurations *A* of *X*_|*V* |_. The paths can be again reconstructed by keeping track of which configuration *B* was used to compute each score in matrix *W*.

The above formulas are not convenient for implementation because we need to iterate over multiple configurations *B* ⊆ *X*_*i*−1_ for each configuration *A* ⊆ *X*_*i*_. It is easier to organize computation in a forward fashion, where we first initialize *W* (*i, A*) to −*∞* for all *A* and then iterate over all configurations *B* of *X*_*i*−1_ and use *W* (*i* − 1, *B*) to update up to *w* + 2 relevant values of *W* (*i, A*), as shown in Algorithm 1. The algorithm clearly works in *O*(2^*w*^ ·*w*· |*V*|) time, provided that sets *A* and *B* can be manipulated in *O*(1) time, which is reasonable since they are used to address the matrix and thus presumably fit into a single computer word.

##### Algorithm 1

Computation of matrix *W* given an incremental path decomposition *X*_1_, …, *X*_|*V* |_ and incremental vertices *v*_1_, …, *v*_*n*_.

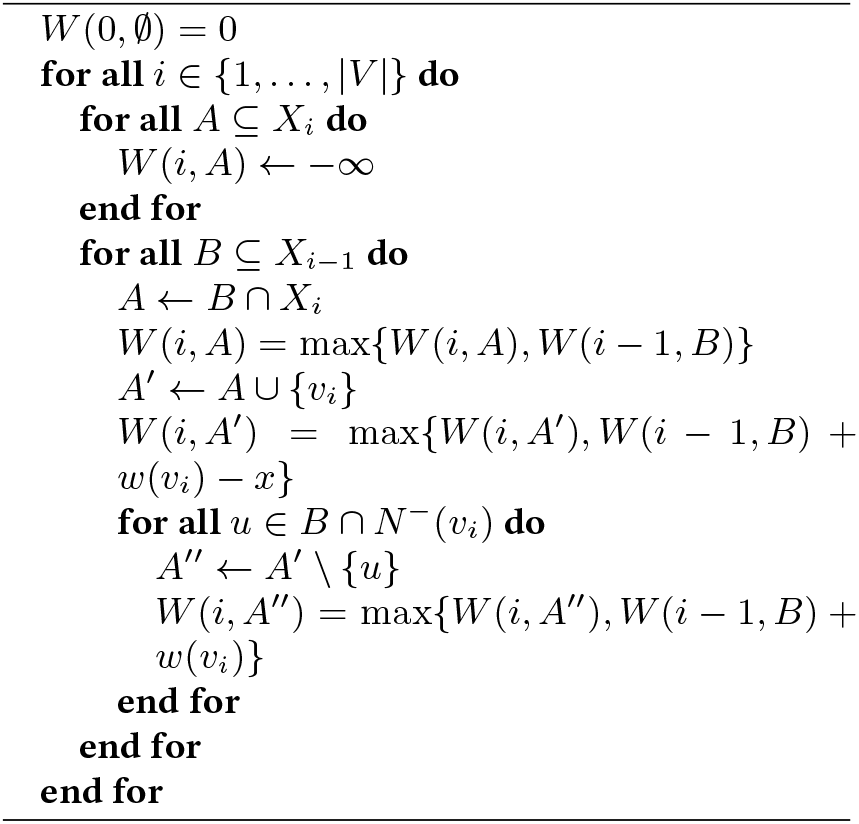

#### Creating an incremental path decomposition

Our algorithm gets the incremental path decomposition as an input. For completeness we describe a heuristic algorithm for creating an incremental path decomposition for a DAG *G*, although, not necessarily the one with the smallest width. Let *v*_1_, …, *v*_*n*_ be a topological ordering of *G*. Put these vertices into subsequent bags, i.e. *X*_*i*_ = {*v*_*i*_}. These bags already fulfill property (*iii*) from Definition 4.1. From Lemma 4.1 we know that the bag where a vertex *v* appears for the first time also contains all *v*’s predecessors. In our case, in each bag *X*_*i*_ a new vertex *v*_*i*_ appears, therefore, add all predecessors of *v*_*i*_ into bag *X*_*i*_. This does not break property (*iii*) from Definition 4.1 and it fulfills property (*i*). To fulfill property (*ii*) in Definition 4.1, find the first and last occurrence of each vertex *v* in the bags, and add vertex *v* into the bags in-between. This does not break property (*i*) and (*iii*) from Definition 4.1 and it fulfills property (*ii*). The complexity of this algorithm is *O*(*w* ·|*V*|).

The resulting path decomposition is incremental. Namely, bag *X*_*i*_ is the first bag where vertex *v*_*i*_ appears, and therefore, |*X*_*i*_ *∖X*_*i*−1_ |*≥* 1. The difference of the sets cannot be 2 or more, as then the other additional vertex has to be *v*_*j*_ where *j* < *i* which means it appeared already in bag *X*_*j*_, and due to condition (*ii*) from Definition 4.1 it means *v*_*j*_ ∈ *X*_*i*−1_.

## 5. Experiments

We created a prototype implementation of the algorithm for *n*-layered bubble graphs from Section 3; this implementation can be found at https://github.com/evicy/thesis. We tested our implementation on the task of identifying GC-rich regions in a pangenome of *Escherichia coli* bacterium. The GC content of DNA sequences, i.e. the percentage of guanine (G) and cytosine (C) bases, is a frequently used statistic when analysing genomes. It has been well studied across organisms, revealing connections between the GC content and various genomic characteristics [20]. GC-rich regions were also used in the study of the maximum segment sum problem by Csűrös [4].

To prepare our data set, we used the complete genome of *E. coli K12-MG1655* as the reference genome [21] and sequencing reads from several strains of *E. coli* isolated from supermarket produce [22]. The reads were downloaded from project PRJNA563564 in the European Nucleotide Archive (ENA) database [23]. We mapped the reads to the genome using BWA [24], processed alignments by SAMtools [25] and then discovered sequence variants for individual strains compared to the reference genome using Freebayes [26], The resulting VCF file with sequence variants was used to construct a elastic-degenerate string by the EDSO [27] tool. Our tool then transforms this EDS to an *n*-layered bubble graph where vertices are single bases (as in Figure 2) and runs our algorithm.

We have tested nine pangenomes, with the first one containing only the reference genome and each successive pangenome adding one additional strain of *E. coli*.

The samples used and the sizes of pangenome graphs are listed in Table 1. To find paths with a high GC content, we assigned weight 1 to bases *G* and *C* and weight -2 to bases *A* and *T*. We tested several values of penalty *x* ∈ {5, 6, 7, 8, 9, 10}. The GC content of E. coli genome is 50.8% on average. These weights mean that the GC content of a selected path is at least 66% to achieve positive score. The penalty ensures that the length of each selected path is at least *x*.

**Table 1:**
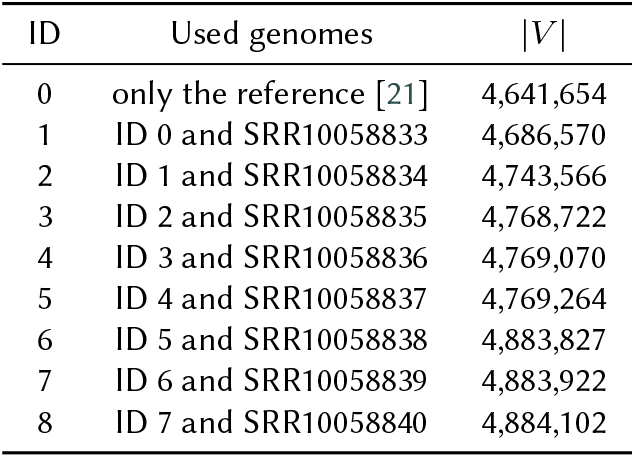
Pangenomes used in our experiment.

In Figure 4, we can see the coverage, i.e. the percentage of the graph that is covered by the selected paths. As expected, the coverages get lower with increasing penalty. By adding genomic sequences to the pangenome, the coverage is increasing, because some of the new variants will introduce *C*’s and *G*’s that can be used by the selected paths.

**Figure 4:**
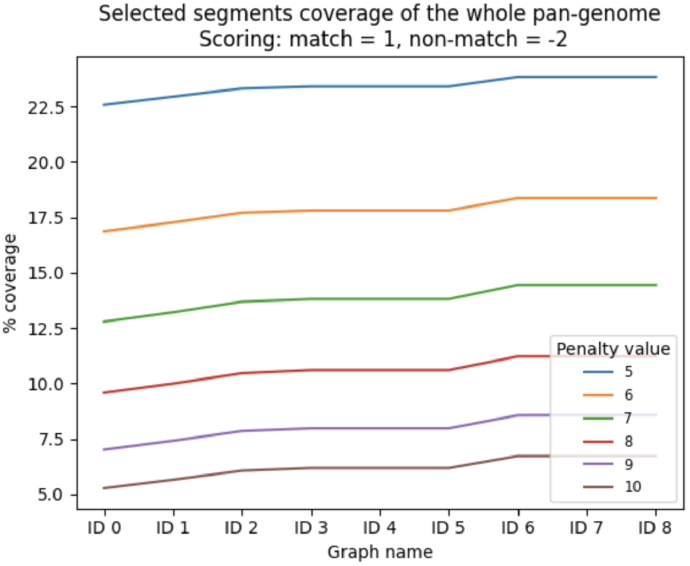
Coverage of a pangenome by selected paths representing high GC clusters.

## 6. Conclusion

In this work, we have defined the maximum-score disjoint paths problem and provided two algorithms for solving it. The first algorithm runs in linear time on *n*-layered bubble graphs, which can represent pangenomes expressed as elastic-degenerate strings. The second algorithm runs on general DAGs in time *O*(2^*w*^ ·*w*· |*V*|) where *w* is the width of a special directed path decomposition defined in this work. We also show the results of a prototype implementation of our first algorithm. In future work, we plan to apply our algorithms to different biological questions stemming from comparative or functional genomics.

Note that our algorithms are purely combinatorial, while many existing approaches for single genomes use statistical methods [28, 10, 29, 30, 31, 32], Csűrös [4] notes that the scores and penalties can be set so that the problem represents finding the maximum likelihood positions of the clusters defined by a two-state hidden Markov model or optimal under complexity penalties, thus providing a link between a combinatorial and statistical versions of the problem. Nonetheless, it is an interesting problem to provide an appropriate extensions of statistical models used in sequence analysis for pangenome graphs.

From a more theoretical point of view, it would be interesting to characterize the complexity of our problem on different classes of directed graphs besides the two studied in this work.

## Acknowledgments

This research was supported by the Operational Program Integrated Infrastructure project ITMS:313011ATL7 co-financed by the European Regional Development Fund. The research was also supported by a grant from VEGA 1/0463/20 and the European Union’s Horizon 2020 research and innovation program (PANGAIA project #872539 and ALPACA project #956229).

